# Retrotransposon LINE1 trans-inhibiting differentiation maintains embryonic stem cell identity

**DOI:** 10.1101/2024.12.02.626147

**Authors:** Likun Ren, Yue Teng, Wenqiu Xu, Jingya Hou, Mengyue Lu, Feitong Shi, Jie Ren, Caihong Zheng, Jun Cai

## Abstract

Long Interspersed Nuclear Elements-1s (LINE1s) are one of the >17% most abundant Retrotransposons in mammalian genomes. LINE1 RNA is high expression during early embryonic development, meanwhile is under tight epigenetic control. Some studies have confirmed LINE1 play a critical role in early embryo development and ESC identity. Previous studies focused on the role of LINE1 in cis-regulation, however, the function of L1 RNA in tans-regulation remains largely unknown. In this study, we performed a high-throughput proximity MARGI (pxMARGI) experiment found transposon LINE1 RNA trans-regulated prefer to young LINE1 subfamily, and in hybrid and non-hybrid two different manner repress ESC to dual progress of 2C-like cells and differentiation. In differentiation process, LINE1 RNA as a scaffold recruit polycomb core subunits combine three core pluripotent factors maintain ESC identity. In 2C process, LINE1 RNA in a sequence-specific manner recruit Kap1 to old L1 subfamily, and recruit ELL3 to RE 5‘UTR maintain ESC self-renewal. Our data point to LINE1 transcriptions more detail trans-regulation mechanism to orchestrate developmental progression for the self-renewal of embryonic stem cells (ESCs). LINE1 RNA may be potential biomarker for noninvasive embryo selection.

## Introduction

Through independent replication of their sequences, transposable elements (TEs) become a group of the most abundant genetic materials within the genome. They account for approximately 46% of the human genome and 37.5% of mouse genome, respectively (Mouse Genome Sequencing et al. 2002). Long Interspersed Nuclear Elements-1s (LINE1s) are one of the >17% most abundant retrotransposons in mammalian genomes (Lander et al. 2001; Mouse Genome Sequencing et al. 2002). Extreme minority possess autonomous retro-transposition, representing a threat to genome stability (Mita and Boeke 2016), whose expression or mobility could be silenced in cells by some epigenetic mechanisms, such as DNA methylation (Deniz et al. 2019) and histone modification (Bulut-Karslioglu et al. 2014; Imbeault et al. 2017) and other mechanisms (Pezic et al. 2014; Pizarro and Cristofari 2016; Warkocki et al. 2018), but many regulatory mechanisms in which are still unclear.

Retrotransposon LINE1 is high expression during early embryonic development, meanwhile is under tight epigenetic control (Fadloun et al. 2013; Low et al. 2021). In recent studies, LINE1 transcription leads to global chromatin opening in mouse early embryos (Jachowicz et al. 2017). LINE1 RNA knockdown cannot recruit Nucleolin and Kap1 to repress transcription factor DUX, then make mESC progress to two-cell (2C) embryo, and regulate genome compartmental organization (Percharde et al. 2018). In naïve ESC, ELL3 occupies at the 5’ untranslated regions (UTRs) of L1Md_Ts, through regulating the activity of the L1Md_T-based enhancers silencing naïve key pluripotency factors (Meng et al. 2023). LINE1 5’UTR as enhancers can physically contact their distal cognate genes, and actives long–range gene expression (Li et al. 2024). Increasingly, researchers have realized the importance of these transposable DNA sequences as a potent source of cis-regulatory DNA motifs to recruit specific regulatory factors, regulate the functional activities of coding genes (Bourque et al. 2008; Kunarso et al. 2010; Testori et al. 2012; Sundaram et al. 2014) and establish the 3D chromatin structure (Lu et al. 2021). And they function as regulatory lncRNAs in ESC self-renewal and in neuronal differentiation (Mangoni et al. 2023). Besides, partly due to technical limitations, previous studies focused on the role of LINE1 in cis-regulation, however, the function of LINE1 RNA in tans-regulation remains largely unknown.

The current development of advanced large-scale experimental biotechnologies, such as MARGI technology (Sridhar et al. 2017), GRID-seq (Li et al. 2017), ChAR-seq (Bell et al. 2018), and RADICL-seq (Bonetti et al. 2020), enables the whole-transcriptome profiling of RNA-chromatin interactions. These high-throughput experimental data provide a special opportunity for us to explore the interactions between transposon-derived RNAs and chromatin for the understanding of the potential trans-regulatory function of these transposon RNAs. Researchers initiated to utilize the RNA-chromatin interaction data for the discovery that transposon RNAs modulate the process of heterochromatin organization (Hao et al. 2020; Chen et al. 2021; Liu et al. 2021).

In this study, we performed a high-throughput proximity MARGI (pxMARGI) experiment in E14Tg2a cells and interpreted the roles of transposon RNAs from the perspective of RNA trans-regulation on gene transcription in embryonic stem cells (ESCs). We reported that transposon LINE1 RNA trans-regulated prefer to young LINE1 subfamily, and in hybrid and non-hybrid two different manner repress ESC to dual progress of 2C-like cells and differentiation. In differentiation process, LINE1 RNA as a scaffold recruit polycomb core subunits combine three core pluripotent factors maintain ESC identity. In 2C process, LINE1 RNA in a sequence-specific manner recruit Kap1 to old L1 subfamily, and recruit ELL3 to repeat element (RE) 5’UTR maintain ESC self-renewal. Our data point to LINE1 transcriptions more detail trans-regulation mechanism to orchestrate developmental progression for the self-renewal of embryonic stem cells (ESCs).

### 1. Transposon RNAs interacting with chromatins has subfamily preference

To systematically investigate characterize and function of repeat RNA (reRNA), which interact with their genomic at the whole genome. We performed a high-throughput pxMARGI experiment in mouse E14Tg2a cells for analysis. The results were demonstrated is very efficiency (Figures S1A-S1D). Further, combined with the public of human H9 MARGI data for analysis, we used effective and rigorous criteria to define the distal reRNA-DNA chromatin interaction, clearly distinguish from the proximal interaction. The analysis indicated that distal interactions of E14Tg2a pxMARGI exhibited high reproducibility and correlation (Figure 1A). Here we have the data from that distal interaction between reRNA and their chromatin DNA, in which about 91.28% interactions are inter-chromosome (Figure 1A-1B). Previous research reports, transposable element (TE) domains located in lncRNA play an important function as DNA-binding regulatory domains. While a growing number of TE insertions in lncRNAs have been experimentally defined as regulatory domains (Johnson and Guigo 2014), the answer is yet unknown how many such TE domains in expressed lncRNAs indeed act as DNA-binding regulatory domains. To know the answer, we integrated analysis reRNA-chromatin and annotated lncRNA database. In human H9 cells, we find that 95% of lncRNA, which have repeat RNA insertion domains, were included reRNA-chromatin distal interaction data. But these repeat regulatory domains account for less than 8% of the total reRNA-chromatin interaction data from pxMARGI (Figure 1C). In mouse E14 cells, about 92% of TE insertion domains in lncRNAs have the role of DNA binding, accounting for only 4% of the total reRNA-chromatin interaction from pxMARGI (Figure 1C). These results suggest that trans-regulation of transposon RNAs on chromosomal DNAs far outweighs the effect of repeat regulatory domains in lncRNAs. Further analysis revealed many kinds of repeat elements, such as LINE, LTR, and SINE, as potential trans-regulating RNAs interactions with chromosomal DNAs (Figure S1E), the conserved LINE and LTR in humans and mouse, in which the LINE-1 family in LINE more prefer to interact with chromatins (Figure 1D). Meanwhile, in order to further confirm this distal interaction between LINE1 RNA and chromatin, we also conducted pxMARGI experiments on LINE1 RNA knockdown E14Tg2a (L1KD), and the results confirmed that after knocking down LINE1 RNA, the trans-regulating interaction between LINE1 RNA and chromatins was significantly reduced (Figure 1E, S1F-G). Therefore, we speculate that LINE1 RNA plays an important trans-regulatory role in the chromatin binding region. To analysis in more detail the LINE-1 subfamily interactions with chromatins. First, we considered the fact that various transposon RNAs have different expressed copies in different species or cell types, we normalized the chromatin-interacting intensities of subfamilies transposon RNAs to ensure removal of the effect of RNA copies. The normalized chromatin-interacting intensity of LINE-1 subfamily revealed that stronger chromatin-associated interaction occurs in the corresponding subfamily of L1PA3, L1PA4, L1PA5 or L1PA7 in human or L1Md in mouse (Figure 1F).Then, when we incorporate the evolutionary ages of TEs (Giordano et al. 2007), it is found that the LINE-1 subfamilies interacting with chromatins in human and mouse cells are concentrated in the younger branches of L1PA and L1Md (Figure 1F).

**Figure 1.**
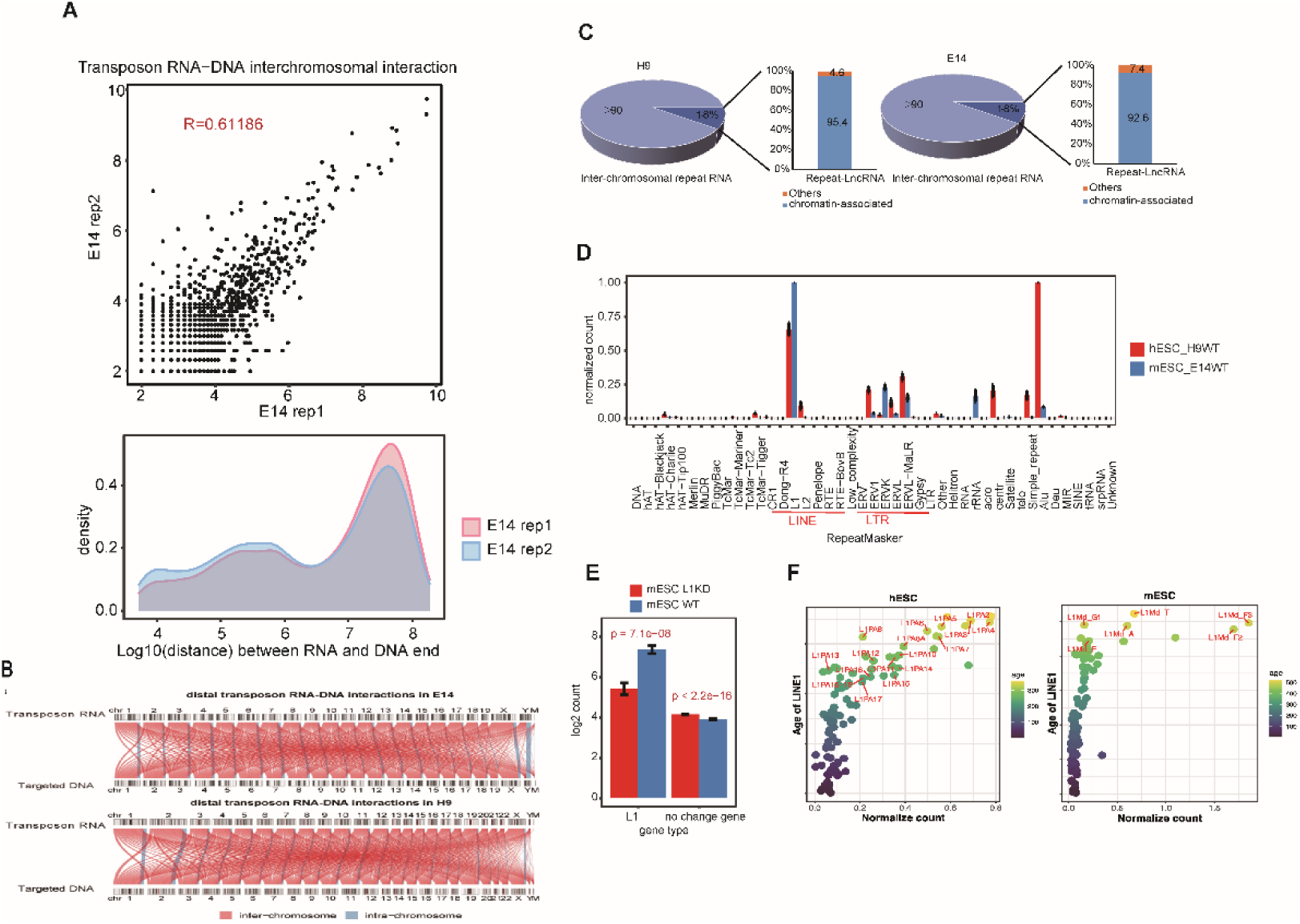
LINE1 RNA trans-regulated chromatin prefer to younger LINE1. (A) The correlation of transposon RNA-DNA inter-chromosomal interactions detected in E14Tg2a pxMARGI replicates (Upper graph). The distribution of the coordinate distance between RNA and paired DNA in each inter-chromosome (Lower graph). (B) The connection map of distal repeat RNA-chromatin interactions across chromosomes in E14Tg2a, H9. Each line represents a group of intra-chromosome (blue lines) or inter-chromosome (red lines) repeat RNA-chromatin interaction events between a pair of chromosomes. The thickness of the line indicates the number of interactions. (C) The proportion of Inter-chromosomal repeat RNA in Repeat-LncRNA of E14Tg2a and H9 (D) The CPM (counts per million mapped reads) of repeat classes using filtered RNA ends that are fully within specific repeat classes and from inter-chromosome. Experiments were independent replicated for two times. (E) The CMP of LINE1 RNA interaction with chromosome between wild type (WT) and knockdown-line1 E14Tg2a. (F) The scatter plots represents enrichment and expression of LINE-1 RNA subfamilies derived from identified inter-chromosome interactions in E14Tg2a, H9 cells. The red labels represents outliers based on cook’s distance using the traditional threshold of 4/sample size. The color of points represented the age and specific evolutionary branches of LINE-1 subfamilies.

### 2. LINE1 RNA knockdown causes mESC to dual progress of 2C-like cells and differentiation

To reveal a more accurate trans-regulatory role of LINE1 RNA in mESC. We knockdown LINE1 RNA by morpholino antisense oligo (ASO), that caused several results, including two-cell (2C) marker genes expression increased and induction of capacity to form embryoid bodies (EBs) impairment similar to previous reports (Figures S2A-S2C) (Percharde et al. 2018). Then to performed 10× Genomics sequencing to WT and L1KD cells, respectively. All detected 22148 cells, 20558 cells were gets after QC. An average of 3337 genes were detected in each individual cell (Figure S2D). We performed canonical correlation analysis (CCA) to normalize variance and correct batch effects between two samples. The results indicated six cell clusters featured by the expression of known marker genes were annotated, including Pluripotent stem cell, Extraembryonic ectoderm cells (ExE), Apoptosis cells, Mesoderm cells, 2C-like cells, Endodermal cells (Figure 2A). Using multiple public data shows a high consistency between our defined cell types and the public data (Figure 2B). When compared with embryonic stem cells, E6.5-E8.5 Epiblast and Primitive streak, result shows Pluripotent stem cells had the highest similarity (Pijuan-Sala et al. 2019), while Gm21761+ESC cells similar with 2C−like cells (Fu et al. 2020). Then the 2C-Like cell our defined was most similar to the 2C-Like cell induced by zhangyi et al (Fu et al. 2020). Compare to two E6.5 public data, we found that ExE ectoderm have the highest similarity in ExE endoderm, ExE ectoderm of E6.5-8.5 (Pijuan-Sala et al. 2019), and ExE ectoderm of E6.5-7.75 (Scialdone et al. 2016); Mesoderm cells had the highest similarity to Mesenchyme and Cardiomyocyte cells in E6.5-8.5 (Pijuan-Sala et al. 2019), and similar to Extraembryonic mesenchymal stem cells in Guo (Wang et al. 2023). Endodermal cells had the highest similarity with Visceral endoderm in E6.5-8.5 and E6.5-7.75 data (Scialdone et al. 2016; Pijuan-Sala et al. 2019) (Figure 2B). Each marker gene significantly high expression in corresponding cluster. For example, 2C-Like cells expresses greater levels of Zscan4d and Zscan4f; Extraembryonic ectoderm cells expresses greater levels of Mt1 and Mt2; Endodermal cells expresses greater levels of Gata6 and Sox17; Mesoderm cells expresses greater levels of Krt8 and Krt18 (Figure 2C). Further analysis showed that the cell numbers of differential clusters were changed when LINE-1 RNA knockdown (Figure 2D). By comparing the cell proportion and cell ratio between WT and L1KD, the results showed that embryonic stem cells and endoderm cells decreased significantly after LINE-1 RNA knockdown, while the number of 2C-like cells, Extraembryonic ectoderm cells and Mesoderm cells increased (Figure 2D). This indicates that LINE-1 RNA knockdown can promote the generation of 2C-Like cells, and the multi-direction progression of ESC to mesoderm and extra-embryonic cells. In order to confirm L1KD induced to different process, first we performed RT-qPCR, results showed increased expression of several differentiation-related genes (Figure S2E). And then for evidence the dual progress at the same time, we performed RNA fluorescence in situ hybridization (RNA-FISH) experiment with Endoderm markers Gata6 and Sox17, 2C-Like cell markers Zscan4d and Zsan4f. The results showed that in the same dish of L1KD cells, Gata6 and Zscan4d (Figure 2E), or Sox17 and Zscan4f (Figure S2F) exist simultaneously. All the above results demonstrated that LINE1 RNA knockdown generated mESC to 2C-Like and differentiation dual process. Furthermore, we also found a cluster of cells with low UMI, and then carried out GO function analysis on genes of the cluster, and observed that apoptosis related functional terms were highly enriched (Figure 2F). Then the mitochondrial proportion, nUMI and nGene contents of different cell types were analyzed. It was observed that the mitochondrial contents of apoptotic cell cluster was significantly higher than those of other cell clusters, while the nUMI and nGene contents were significantly lower than others (Figure 2G). It is consistent with a significantly reduced in ESC proliferation after LINE1 RNA knockdown (Figure S2C).

**Figure 2.**
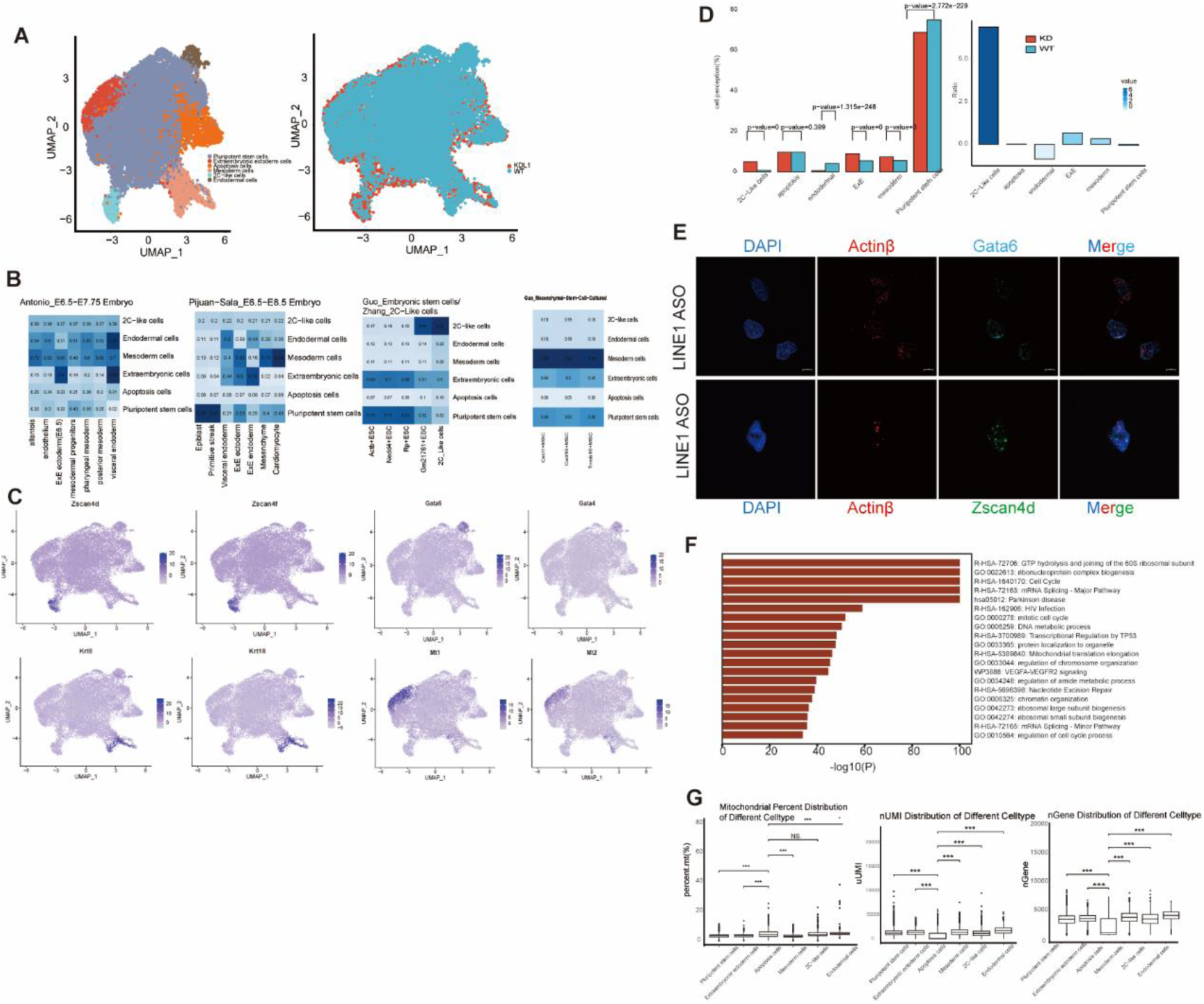
LINE1 RNA knockdown causes mESC to dual progress of 2C-like cells and differentiation. (A)10× genomic single-cell RNA-seq data of WT and L1KD E14Tg2a indicate exist two progress. Two-dimensional projection (uniform manifold approximation and projection (UMAP)) representation of six cell types within two groups (left). Correct batch effects between two samples (right). (B) Hierarchical clustering of marker genes for differential cell types in mouse early embryos or mESC. Heat map showing pairwise Pearson’s correlation coefficients between different public data identified cell types. (C) The expression of marker genes in each cell cluster. (D) The cell proportion and cell ratio have changed when LINE1 knockdown in mESC. (E) RNA-FISH indicated in the same dish of L1KD cells, Endoderm markers Gata6 and Sox17, 2C-Like cell markers Zscan4d and Zsan4f exist simultaneously. (F) GO function analysis of a cluster of cells with low UMI. (G)The analysis of the mitochondrial proportion, nUMI and nGene contents in different cell types.

### 3. LINE1 RNA as a scaffold regulates mESC differentiation process

To investigate the regulatory mechanism of LINE1 RNA in differentiation progress. We integrated E14Tg2a L1KD MARGI data and publicly L1KD RNA-seq data to analysis, 213 genes (34.86%) were detected in 611 up-regulation genes. These genes were called LINE1 RNA trans-regulating genes, LRTGs. Gene ontology (GO) analysis of LRTGs revealed an extremely significant enrichment to nervous and development related hierarchies, including nervous system development, animal organ morphogenesis, cell fate commitment, negative regulation of cell proliferation, cell development, among others (Figure 3A). Certain key genes of those have been reported closely with embryo development and differentiation. For example, Fosl1, Fgf5, Bml1 (Figure S3A). LRTGs binding chromatin region is mainly distributes in gene promoter (-5kb∼+1kb), intron and intergenic regions (Figure S3B). The binding of promoter regions suggests that these interactions may be involved in gene expression regulation.

**Figure 3.**
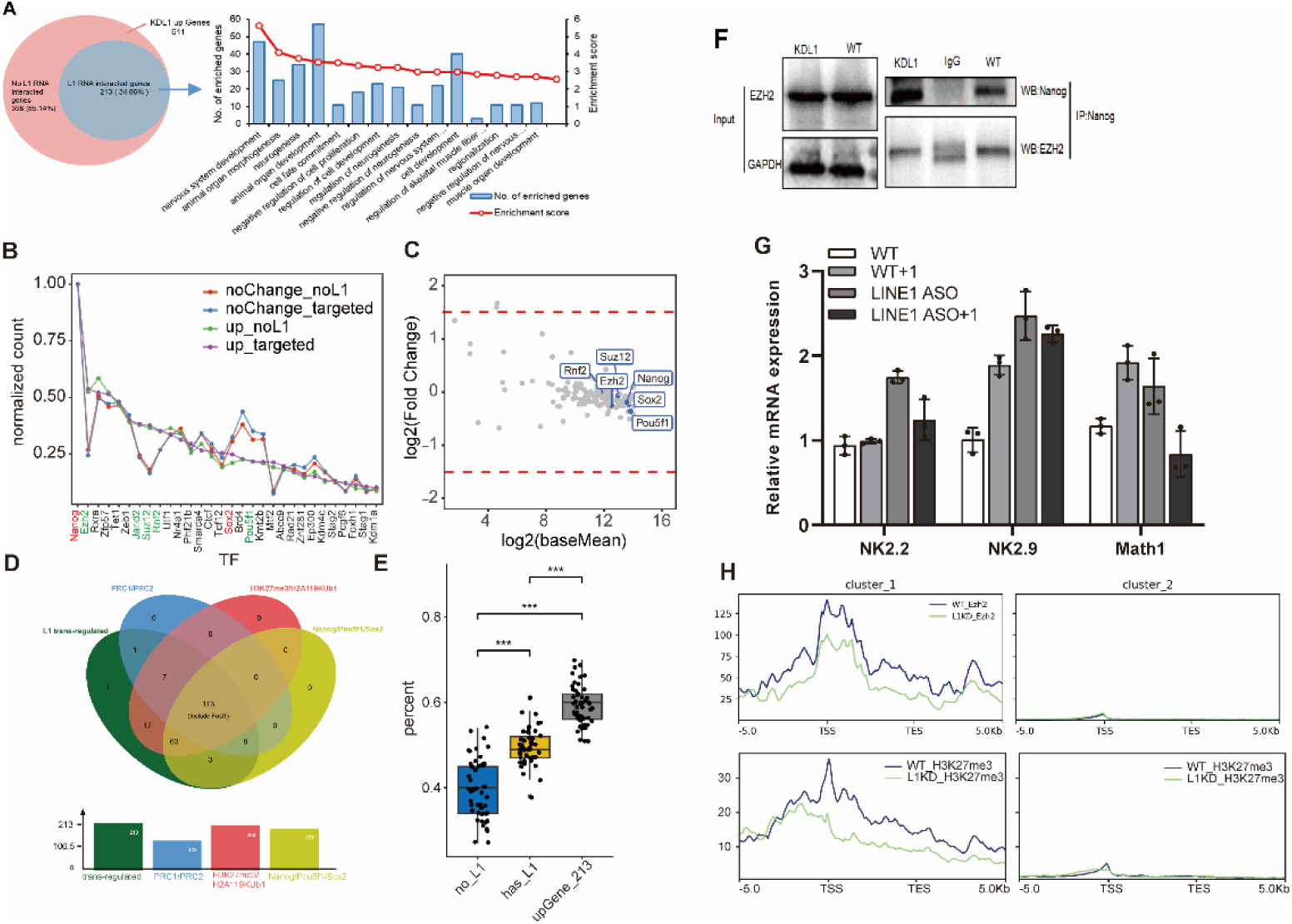
LINE1 RNA as a scaffold combine with Nanog and Ezh2 regulates mESC differentiation process. (A) Gene ontology (GO) analysis of LRTGs (LINE1 RNA trans-regulating genes). (B) The sort of transcription factors co-bound with LINE-1 RNA in the LRTGs promoter regions. The binding sites of transcription factors were annotated by ChIP-seq data in GTRD (http://gtrd.biouml.org). (C) The expression changes of key transcription factors with LINE-1 RNAs in the promoter regions of the LRTGs after LINE-1 knockdown. (D) The Venn plots represent the overlap of co-binding by at least one PRC core subunit, core pluripotent factors, repressed histone modifications and KDL1 up-regulated genes. (E) LINE-1-associated PRC binding chromatin regions were significantly co-located with three core pluripotent factors regions, compared with the PRC-bound regions alone, especially the 213 LRTGs promoter regions in the promoter regions. (F) Co-immunoprecipitation (coIP) demonstrated that Nanog interact with Ezh2 in E14Tg2a, also the interaction intensity was reduced when LINE1 RNA knockdown. (G) Knockdown LINE1 RNA led to upregulation a panel of PRC-repressed genes binding with LINE1 RNA, NK2.2, NK2.9 and Math1. Not change in L1KD. (H) Knockdown LINE1 RNA caused both Ezh2 and H3K27me3 enrichment were decreased in L1KD upGenes.

We using the most comprehensive publicly protein ChIP-seq data from Gene Transcription Regulation Database (GTRD) to infer the transcription factors potentially involved in the regulation of LRTGs in E14Tg2a. First, sorted transcription factors according to the degree of binding in promoter regions, two types of transcription factors were found to be more co-bound with LINE-1 RNA in the LRTGs promoter regions, including maintaining pluripotency factors in embryonic stem cell Nanog and the Polycomb Repressive Complex (PRC) core components Ezh2, Suz12, Rnf2, and the accessory subunit Jarid2 (Figure 3B).The expression of transcription factors were altered after knockdown by exclude (Figure 3C), meanwhile RT-qPCR results showed same results, but Nanog was increased expression (Figure S3C). Future Based on ChIP-seq data, we also found that these regions binding by at least one PRC core subunit were also binding by maintaining pluripotent factors, co-binding regions about 93.8% (121/129) (Figure 3D). In addition, in the promoters of all genes, we found that the PRC and LINE-1-associated PRC binding chromatin regions were significantly co-located with three core pluripotent factors regions, compared with the PRC-bound regions alone, especially the 213 LRTGs promoter regions (Figure 3E). Therefore, we inferred that the presence of LINE1 RNA may promote the co-location of Polycomb repression complex core subunits and pluripotent factors in the LRTGs promoter region. To validate the finding we perform Co-immunoprecipitation (coIP), results demonstrated that Nanog was able to interact with Ezh2 in E14Tg2a, also the interaction intensity was reduced when LINE1 RNA knockdown (Figure 3F). Next, we attempted to further confirm the LINE1 dependency between Pluripotent factors and PRC core subunits. To this end, we treated KDL1 and WT with Ezh2-special inhibitor, Tazemetostat (Ding et al. 2023), results indicated that knockdown LINE1 RNA led to upregulation a panel of PRC-repressed genes, NK2.2, NK2.9 and Math1, which were LINE-1 RNA binding in WT. Nevertheless, not change in L1KD (Figure 3G), indicated LINE-1 RNA knockdown led to PRC core subunits fall-off at the same time. In addition, knockdown LINE-1 RNA induced increased expression of inhibitory developmental genes co-located by LINE-1 RNA, PRC and pluripotent factors (Figure S3C), such as Fosl1. Ezh2 is an important catalytic enzyme of the Polycomb Repressive Complex 2 (PRC2) complex, which is responsible for histone modifications of H3K27me3 that silence the expression of developmental regulator genes. To focus on these developmental genes, we found that knockdown LINE1 RNA caused both Ezh2 and H3K27me3 enrichment were decreased (Figure 3H), but enrichment of pluripotent factors did not change (Figure S3D). Together, these results indicated that LINE-1 RNA might as a scaffold combine with Nanog, then specifically recruit Ezh2 to Nanog-LINE1 complex regulate development regulator genes.

### 4. Optional binding modes between transposon RNAs and chromatin regions

Next, we wanted to know how way LINE1 RNA interaction with chromatins. First of all, we analyzed the distribution of the DNA end of MARGI data on the genome and found that the chromatin regions bound by the repeat LINE-1 in E14Tg2a were significantly derived from the same family of repeat elements (Figure 4A), suggest reRNA-chromatin interaction maybe in a sequence-mediated manner. Next, Predicated RNA-DNA Watson-Crick complementary duplex and triple-helix structures formed by the sequence context of the paired transposon RNA and its targeting DNA region with software (Kruger and Rehmsmeier 2006) DNA sequences outside the transposon RNA targeting regions were randomly sampled to compose the pseudo transposon RNA-DNA pairs as control, and the distal interaction pairs between non-transposon RNAs and chromatins were selected as another control group. The distribution of minimal free energy defining the binding strength of RNA-DNA hybrid duplexes in the sequence contexts of the paired ERV, rRNA or LINE-1 RNA and its targeting DNA region has the unique characteristics of long tail and bimodal (Figure 4B, Figure S4A). These ERV, rRNA or LINE-1 RNA-chromatin interactions form RNA-DNA hybrids with ultra-low minimal free energy less than -60 kcal/mol, the binding strength of which is competitive with that of the RNA-DNA hybrids formed by pairwise perfect-matched sequences (Figure 4B, Figure S4A). The binding strength of LINE-1 decreased in L1KD (Figure 4B), but rRNA and ERV increased (Figure S4A-S4B). These results indicate that LINE1 RNA interacts with chromatin in a RNA-DNA hybrid duplexes manner. Few triplex-helix structures, on the contrary, occur in the sequence contexts of transposon RNA-chromatin interactions by predication. But LINE-1 RNA identified in mouse and human embryonic stem cells, which occupied chromatin regions co-location with pluripotent factors, H3K27me3 and PRC were predicted using Triplexator and RNAhybrid (Kruger and Rehmsmeier 2006) software. The results showed LINE-1 RNA mainly by trans-action base on protein, was non RNA-DNA hybrid mode (Figure 4C).

**Figure 4.**
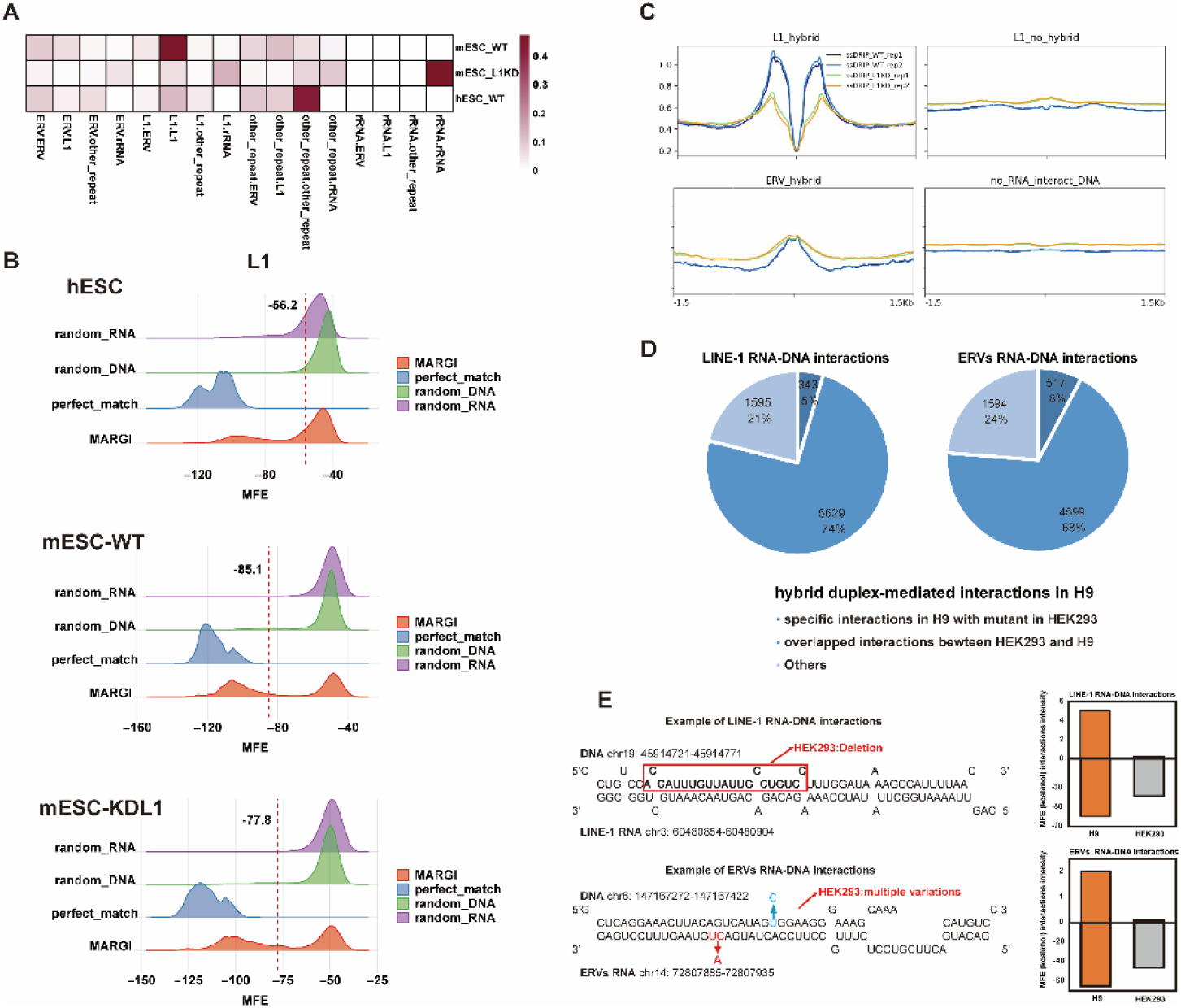
RNA hybrid predicts the intensity of interactions between repeat RNAs and Chromatin. (A) Heatmap showed the relative proportion of our defined confident DNA loci targeted by LINE-1, ERVs, rRNA and RNA outside of annotated repeat elements according to whether overlapped repeat elements and which repeat family they overlapped. (B) showed MFE density of RNA hybrid prediction from selected repeat RNA-DNA or non-repeat RNA-DNA read pairs. Background data was randomly sampled 10,000 150 bp DNA loci in whole genome and combined with randomly sampled repeat RNA from defined confident read-pairs for 100 times. Perfect match was positive duplex-mediated interactions. Their RNA hybrid result was perfect matched allowing G:U wobble base pairs in defined confident interactions, respectively. Purple dotted line is 95 % confidence interval in randomly sampling. (C) ssDRIP-seq experiment indicated that the signal of near the LINE1 RNA hybrid region are significantly reduce in L1KD. (D) Based on duplex-mediated LINE-1 or ERVs RNA interacted DNA loci in H9, the pie plots showed the proportion of common identified in H9 and HEK293 cells and the influence of HEK293 genome variations on identified LINE-1 or ERVs RNA-DNA interactions. Specific examples of the influence of HEK293 genome variations are showed in (E).

Stable RNA:DNA hybrid structure is usually accompanied by the unwinding of double-stranded DNA and the formation of R-loop. Therefore, we conducted the ssDRIP-seq experiment (Figure S4C), and the results confirmed that the signal of near the LINE1 RNA hybrid region was significantly reduced in L1KD (Figure 4D). Further analysis showed that the LINE1 RNA forming hybrid was older than that forming no-hybrid (Figure S4D).

Comparisons between different human cell types demonstrate that about 70% hybrid-mediated LINE-1 or ERV RNA-chromatin interactions are consistent in H9 and differentiated HEK293 cells, free from cell identity. Genomic variations (Lin et al. 2014) on the RNA-DNA hybrid duplexes abolish about 5% transposon RNA-chromatin interacting activity between H9 and HEK293 cells, as described in the following two specific examples (Figure 4E). A stable RNA-DNA hybrid duplex with minimal free energy of -60.5 kcal/mol is formed that mediates the interaction between a LINE-1 RNA and a piece of DNA sequence on chromosome 19q13 in H9 cells. An 18bp deletion in the DNA sequence of this RNA-DNA hybrid in HEK293 cells leads to the reduction of the RNA-DNA binding strength with minimal free energy of -38.2 kcal/mol (Figure 4F). Meanwhile, in HEK293 cells the LINE-1 RNA-chromatin interaction is no longer detected at this region with the MARGI experiment data, although there are no differences between H9 and HEK293 cells in the expressions of other potential trans or cis-regulatory factors around this region. Similarly, another case, where multiple single nucleotide variations occur in the ERV RNA and its targeting LUADT1 gene body on the chromosome chr6q24 in HEK293 compared with H9 cells, demonstrates the loss of ERV RNA interaction is result from the weakening of the RNA-DNA hybrid (Figure 4F). All the above evidence reveals that the RNA-DNA hybrid binding mode is required for many transposon RNA-chromatin interactions.

### 5. LINE1 participates in the mESC identity maintain by DNA-RNA hybrid manner

These results above reveal many LINE-1 RNAs interact with chromatins via the mode of RNA-DNA hybrid duplexes (Figure 5A). In order to further explore the regulatory effects of LINE1 RNA on chromatin, first, we screened out which transcription factors were involved by DNA-RNA hybrid manner. These co-binding proteins include Kap1 as well as Zeb1 in mESC, Kap1 and ZNF486 in H9 more dependent on LINE1 RNA sequence information. Kap1 transcription factor is also enriched in the ERV-bound chromosome region (Figure S5A). Previous studies observed that in mESCs, LINE1 acts as a nuclear RNA scaffold recruiting Kap1/Trim28 to repress Dux, which is an activator of the transcriptional program in zygotic genome activation, maintain ESC identity (De Iaco et al. 2017; Percharde et al. 2018). Furthermore, the R-loop signals (Sanz et al. 2016) and the minimal free energy (MFE) of RNA-DNA hybrid support it is the RNA-DNA hybrid duplex in the LINE-1 RNA and chromatin interactions that guides the LINE-1 RNA-Kap1 complex for the recognition on Dux gene (Figure 5B). The protein binding and histone modification data and the PRC-KO experiment data further indicate that PRC acts as an essential partner for the binding and repression on the Dux gene that is co-regulated by the LINE-1 RNA and Kap1 in mESCs (Figure 5C). Besides Dux, the RNA-DNA hybrid duplex mediates the LINE-1 RNA-Kap1 for the binding and inhibition on some other genes relative with ESC differentiation (Figure 5C). Among these genes, Adcy2 plays an important role in embryonic development including cell migration, proliferation, and differentiation (McCallie et al. 2019). And C130026I21Rik, a homolog for the gene SP140 in human, participates in biological process of cell-cell adhesion that is crucial to maintain tissue morphogenesis and regulate cell migration and differentiation during development (Kashef and Franz 2015; Karaky et al. 2018). The high-throughput MARGI data also reveal many distal interactions between LINE-1 RNA and LINE-1 DNA with a co-binding partner of KAP1, significantly reduced in L1KD (Figure 5D). The interactive LINE-1 RNAs and LINE-1 DNAs are dominantly derived from subfamilies of L1PA2, L1PA3 and L1PA4 in human consistent with previous findings that KAP1 (TRIM28) inhibits age-specific LINE-1 lineages in human embryonic stem cells (Castro-Diaz et al. 2014) (Figure 5E). In mouse embryonic stem cells E14 is more concentrated in L1Md_F2 and L1Md_T subfamilies (Figure 5E), and more enriched in younger subfamilies L1Md_A and L1Md_T in L1KD (Figure S5B).

**Figure 5.**
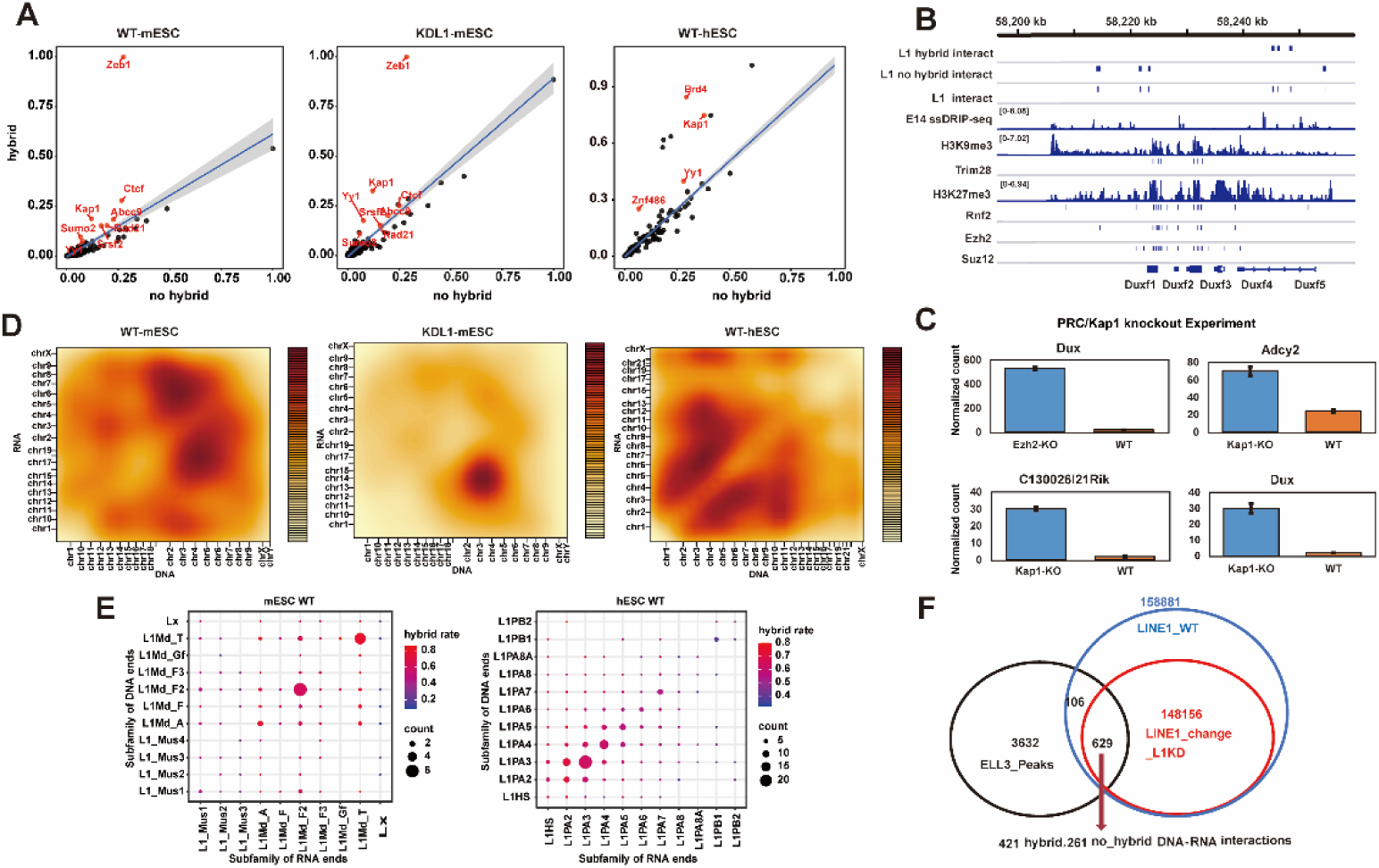
LINE1 as guide RNA interact with chromatin by hybrid manner maintain mESC identity. (A) The number of DNA loci co-localized by LINE1 RNA and the protein downloaded from GTRD. X-and y-axis represent the type of interactions of LINE1 RNA. (B) IGV show the ChIP-seq binding sites of the components of PRC, Kap1 from GTRD, R-loop signals in DRIP-seq, two different ways of LINE1 RNA to binding chromatin location. The coverage of LINE1 RNA and histone modifications related to the above-mentioned proteins around Dux gene. (C) The changes of these genes expression after Kap1 or PRC knockout. (D) The chromosomal coordinate heatmap show the number of distal interactions between LINE1 RNA-Kap1 and LINE1 DNA in E14, KDL1 and H9 cells. The whole genome is in 10Mb/bin resolution. (E) The similarity of subfamilies between LINE1 RNA and trans-interaction LINE1 DNA with a co-binding partner of Kap1.The size and shade of the color represent the rations of RNA:DNA hybrid-mediated distal interactions and the level of similarity, respectively. (F) The overlap of ELL3 ChIP-seq peaks with KDL1 change DNA loci largely are hybrid manner.

Previous research reports 2273 ELL3 chip-seq peaks located in RE, 85% of that overlap L1Md_Ts in mESC. At the same time, MERVL and the 2C genes are upregulated after Ell3 KO or LINE1 ASO (Meng et al. 2023). Based on the above conclusions we inferred that L1Md_Ts induced ELL3 bind to RE in hybrid manner. We identified 735 ELL3 chip-seq peaks overlapped with the distribution of the DNA end of MARGI data on the genome, of which 629 peaks were changed when knockdown LINE1. Therefore, we inferred that 629 regions are regulated by LINE1 RNA and ELL3 together, and further analysis revealed that 421 RNA-DNA interactions pairs of 636 regions are hybrid, 261 RNA-DNA interactions pairs of 636 are no-hybrid. Based on above of the results we conclude that LINE1 RNAs guides ELL3 for DNA loci by RNA-DNA hybrid manners (Figure 5F).

### 6. LINE1 as a potential marker for embryo quality identification

Generally, LINE1 transcription in somatic cells is strictly inhibited by a variety of mechanisms. However, LINE1 is disproportionately high expressed during highly conserved early embryonic and embryonic stem cell (ESC) development (Jachowicz et al. 2017; Percharde et al. 2018; Li et al. 2024). Increasingly researches confirms that LINE1 RNA plays an important role in early embryonic and stem cells. At present, a non-invasive method that can accurately screen with the highest potential embryo is an urgent need for clinical assisted reproductive technology (ART). Cell free RNA (cf-RNA) of medium at different development stage of the embryos can dynamically reflect embryo quality in real time, which is more representative of embryo condition than DNA. We inferred that LINE1 RNA could be potential biomarker in distinguishing the embryos with different quality as an alternative or supplementary approach for subjective morphology criteria. We collected 4 blastocyst medium patients who received intracytoplasmic sperm injection (ICSI), two are high-grade embryos and two are low-grade embryos (Figure 6C). Measuring the reads size distribution of the cf-RNA, it exhibited a peak at 110 bp, which was shorter than that in conventional sample type (150 bp) (Figure 6B). Repeat elements analysis of cf-RNA, found that L1 family take a larger proportion in all mediums, but no significant difference between all mediums (Figure 6D).

**Figure 6.**
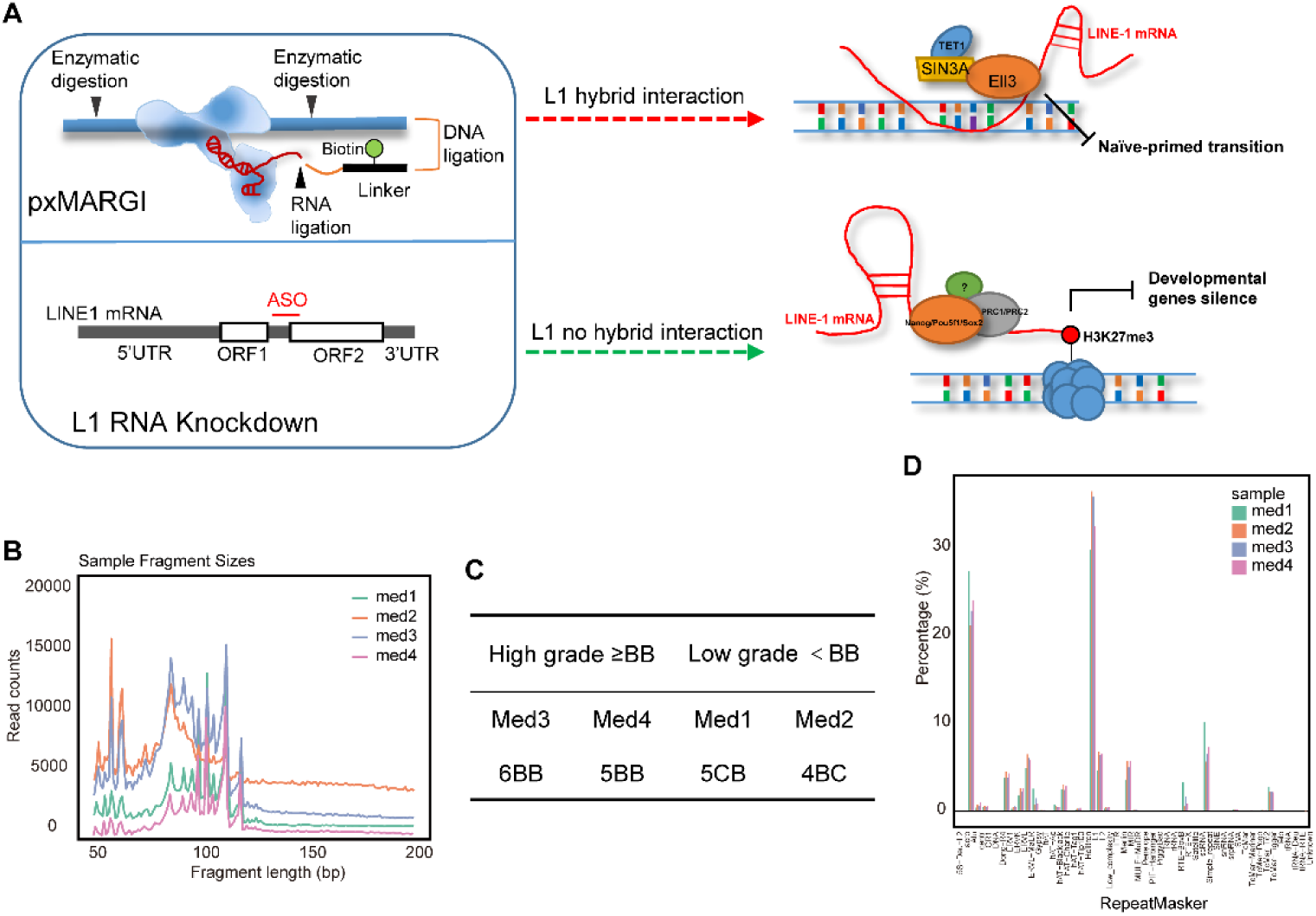
LINE1 is essential for maintain embryonic stem cell identity. (A) Cartoon model illustrating function of LINE1 mRNA in maintain embryonic identity through two different binding modes with DNA. (B) The sequencing reads length distribution in blastocyst medium. Different colors represent different samples. (C) The blastocysts were divided into two groups: high-quality and low quality according to score of blastocyst morphological. (D)The proportion of repeat elements family in different blastocyst medium.

## Discussion

Transposable element play an important role in embryonic stem cells and early embryonic development (Jachowicz et al. 2017; Percharde et al. 2018; Low et al. 2021). LINE1 is highly expressed in stem cells and early embryos and is tightly controlled by a variety of epigenetic modifications (Fadloun et al. 2013; Low et al. 2021). Recently, RNA may be interaction with chromatin via three modes. First mode is cis-interactions, RNA at their sites of synthesis. Second mode is trans-interactions, RNA being released to different genomic loci. Third mode is both cis-and trans-interactions simultaneous effect (Li and Fu 2019). Until now, due to technical reasons, there are relatively few researches on trans-regulation. In recent years, many methods has been developed to identify true chromatin binding RNAs and their interaction sites, each of which has advantages and disadvantages. In particular, the advanced high-throughput experimental biotechnologies exploring RNA-chromatin interactions are enabling the deep understanding of the potential function of transposon RNAs on DNA regulation in cells. When the ultra-short sequencing reads and lower mapping rate of about 20bp reads in GRID-seq (Li et al. 2017) limit the accurate recognition of complex repeat elements, here we use the MARGI technology (Sridhar et al. 2017), which uses a bivalent linker to ligate to RNA and DNA followed by circularization for library construction, with 150bp-length sequencing reads is conducive to the profiling of transposon RNAs and their distal targeting DNA loci. We performed MARGI experiment in mESCs and utilized in-house and public high-throughput data for analysis on trans-regulation of transposon RNAs. And we strictly define the RNA and the site of chromatin as distal trans-regulation DNA loci. It is found that most of distal interactions between reRNA and their chromatin DNA are inter-chromosomal. But MARGI need more sequencing to obtain a sufficient number of mated RNA and DNA reads, acquisition rate of effective data is low. A common drawback of current approaches is modest sensitivity, and, as a result, many low-abundance chromatin-associated RNAs may cannot were detection. Cannot used in precious materials. It is urgent that many new inventions are exploited to study RNA-chromatin interactions.

Our results show that LINE1 RNA as a scaffold recruits Polycomb core subunit Ezh2, promote H3K27me3, inhibits differentiation genes and maintain mESC identity. LINE1 has previously been reported to regulate the development of the cerebral cortex as a long non-coding RNA, a role that appears in E12.5 and is involved in regulating the balance between neuronal progenitor cells and differentiation. In cortical cultured neurons, L1 RNA is primarily associated with chromatin and interacts with the multiple combing inhibition Complex 2 (PRC2) protein subunits Ezh2 and Suz12. According to our study, such effect of LINE1 may appear earlier in the early stage of neural differentiation (Mangoni et al. 2023). After knockdown LINE1 in mESC, we detected the co-existence of ExE endoderm marker genes and 2C-Like genes by RNA-FISH. Thus, we confirm that LINE1 is important for maintaining mESC pluripotency. To date, EZH2 and SUZ12, both are key components of Polycomb repressive complex 2 (PRC2), have been found to have the ability to directly contact RNA (Cifuentes-Rojas et al. 2014; Beltran et al. 2016; Wang et al. 2017). Polycomb is a complex containing many core subunits, and whether it interacts with other subunits at the same time needs further investigation(Di Croce and Helin 2013).

Previous studies have reported the presence of a large number of protein-coding genes and lncRNAs in culture medium, coding genes may serve as potential biomarkers to distinguish different quality embryos, but repeat elements information was not mentioned. The peak length of cf-RNAs reaches 110 bp, which is obviously shorter than 150bp of conventional library in medium, which is reasonable. The machanism is similar to that of cf-RNA in the blood. The large proportion of repeat elements in cf-RNAs can be used to assess the molecular characteristics of medium with different quality, and provide a reference for noninvasive identification and prediction of blastocyst quality. This experiment has many limitations. First, the small sample size restricts the accuracy of the results. Therefore, more clinical studies are needed for further analysis before clinical application. Secondly, cf-RNA is easily degraded, and the low volume medium limits its wide application. In conclusion, the mechanism of cf-RNA needs further study.

## Methods

### Mouse ES cell culture

Mouse E14Tg2A (E14) ES cells were purchased from the National Collection of Authenticated Cell Cultures (Shanghai, China). ES cells were cultured on 0.1% gelatin-coated plates in ES-FBS culture medium (KnockOutTM DMEM, Thermo Fisher Scientific, 10829018), 15% FBS (Biological Industries, 04-002-1A), 0.1mM nonessential amino acids (Millipore,TMS-001-C), 50 U/mL penicillin-streptomycin (Invitrogen, 15140122), 0.1mM 2-Mercaptoethanol (Millipore) and 1,000U/ml LIF supplement (ESGRO, Millipore), EmbryoMax Nucleosides (Millipore, ES-008-D), 2mM GlutaMAX (Thermo Fisher Scientific, 35050061)).

### ASO knockdown LINE1 RNA

ASO knockdown LINE1 RNA was performed according to a previously published protocol (Percharde et al. 2018; Percharde et al. 2021). ASO were synthesized by GENE TOOLS. ASOs were introduced into cells by nucleofection, utilizing an Amaxa Nucleofector 2B device and ES nucleofection kit (Lonza), according to the manufacturer’s instructions. Briefly, prepared 4-5 million ES cells resuspend in 95ul nucleofector solution per sample, and mix with 5ul (5nmol) of ASO. Select program A-023 to nucleofection. And then transfer the cells from the cuvettes to culture plates. These are cells used to other downstream expreiments after 48h post nucleofection.

### Cleavage under targets and tagmentation (CUT&Tag) assay

The CUT&Tag assay was performed using the NovoNGS® CUT&Tag 3.0 High-Sensitivity Kit (NovoProtein, N259-YH01). Collected 2.0 × 105 mESC cells were washed with 200ulL wash buffer, then mixed with 90ul wash buffer and 10ul activated concanavalin beads incubated 10min at room temperature, discard supernatant. Follow by incubations with the primary antibody RT 2h [Ezh2(Cell Signaling Technology, #5246), H3K27me3 (Millipore, 07-449),Out4 (Abcam, ab181557),Nanog (Active Motif, 61419),Sox2 (Cell Signaling Technology, #23064) ],and secondary antibody (RT,1h), the cells/beads complexes were washed twice with antibody buffer, then incubated with pAG-Tn5 for 1 h RT. Then, MgCl2 was added to activate tagmentation for 1 h and washed twice with ChiTag buffer. The tagmentation reaction was stopped by 1ul 10%SDS at 55℃ 10min. The transposed DNA fragments were purified using Tagment DNA Extract Beads and to construct libraries were sequenced using the Illumina HiSeq2000/2500 platform.

### Real-time RT-qPCR

Total RNA was extracted from cells using TRIzol (Sigma-Aldrich, St. Louis, MO, USA). Subsequently, RNA reverse transcription was carried out using the HiScript III RT SuperMix for qPCR (+gDNA wiper) (Vazyme, Nanjing, China) according to the manufacturer’s protocol. The expression of mRNA was analyzed using qRT-PCR with the TB Green® Premix Ex Taq™ II (Tli RNaseH Plus), ROX plus (Takara). Real-time PCR was performed in triplicate using the primers listed in Supplementary Table 1 and a CFX96 Real-time System (Bio-Rad Laboratories, Richmond, VA, USA). Expression levels were normalized to that of glyceraldehyde 3-phosphate dehydrogenase (GAPDH).

### ssDRIP-seq

ssDRIP was performed as described previously, with several modifications. Briefly, 1×107 cells wash one time use PBS, then resuspend in cold cell membrane lysis buffer (5mM Tris pH=7.5; 0.85mM KCl; 0.5% NP40) on ice for 10-20min. PBS wash one time again, then resuspend in TE buffer, followed by adding 10%SDS (final concentration:0.5%) and proteinase K (final concentration:0.1mg/ml), and then incubated in thermo shaker 200rpm at 37℃ for 6h. Add 1/4 volume 5M KAc of mixture, the tubes were mixed and stay on ice for 20min. Then add the same volume phenol: chloroform: isoamylalcohol (25:24:1) into tubes, mixed well and centrifuged 12,000g at 4℃ for 10min. Transferred the supernatant to a new tube,add 1/10 volume of 3M NaAc and 0.7 volume of isopropanol, the DNA was precipitated at -20℃ for 1h. Mixtures were centrifuged 12,000g at 4℃ for 30min. DNA pellet was washed twice with 1ml 75% ethanol, and air-dried, then was dissolve in TE buffer. Take10ug gDNA to a new tube, add 5ul RNase H (M0297,NEB), 20ul RNase H buffer, to total 200ul system, incubated at 37℃ overnight, next day supplement RNase H and buffer, incubated at 37℃ overnight again. The DNA was purified with phenol: chloroform: isoamyl alcohol (25:24:1) extraction as described above.

gDNA was digested with Mse I, Dde I, Alu I, and Mbo I (NEB; final concentration: 100 U/ml for each enzyme) at 37°C for 6h. After purified with phenol: chloroform: isoamyl alcohol (25:24:1) extraction. For each sample, 2ug input DNA was immunoprecipitation with 1× DRIP binding buffer [10mM NaPO4 (pH 7.0), 140mM NaCl, and 0.05% Triton X-100] and 2ul S9.6 antibody (Millipore, MABE1095) at 4°C rotation 10rpm overnight. 100ng gDNA as input for next step. After adding 10ul Dynabeads Protein G (Invitrogen, 10004D) which has been washed with 1× DRIP binding buffer 3 times, for each tube. The beads with DNA/antibody complexes were incubated with rotator at 4℃ for 4h. After binding, beads/DNA/antibody complexes are washed 4 times with 1× DRIP binding buffer at room temperature, each wash takes 10min. Elution buffer [50mM tris-HCl(pH 8.0),10mM EDTA and 0.8 mg/ml proteinase K] was added into mixtures, and the tube was incubated in an Eppendorf ThermoMixer at 55°C for 1h. IP-gDNA were purified with phenol/chloroform extraction as described above, and used for next-generation sequencing (NGS) library construction or qPCR.

ssDRIP-seq library was constructed by using the Accel-NGS 1S Plus DNA Library Kit (Swift Biosciences). Briefly, the DRIPed DNA sample was fragmented to an average size of about 250 base pairs (bp) with an S220 Focused-ultrasonicator (Covaris, Woburn, MA, USA). The sonicated DNA was denatured into ssDNA at 98°C for 2 min and placed on ice immediately for another 2 min. The truncated adapter 1 was ligated to the 3′ end of ssDNA first, followed by extension. The truncated adapter 2 was ligated to the 5′ end. After PCR amplification, libraries were purified with AMPure XP beads (Vazyme).These libraries were sequenced on an Illumina HiSeq2000/2500 platform .

### RNA fluorescence in situ hybridization (RNA-FISH)

ES RNA-FISH was performed using QuantiGene ViewRNA (Affymetrix) based on the manufacturer’s instructions. ES was cultured on ibidi plates during 48h post nucleofection. And then discard medium, PBS wash one time. ES cells directly were fixed in 4% paraformaldehyde (PFA) for 15min at room temperature, were washed three times, 5min per time. Add 200ul detergent solution QC for 5min at RT, PBS washed one time. Digestion with protease for 10min at RT. Followed by Probe sets, preAmplifier Mix, Amplifier and Label probe Mix successively incubated and wash. Last, All nuclei were counterstained with SlowFade Diamond (Invitrogen, S36968). Images of stained cells were acquired by a confocal laser scanning microscopy (Zeiss LSM980). RNA probes are Sox17 (Thermo Fisher Scientific, VB1-10057-VC), ActB_Type1 (Thermo Fisher Scientific, VB6-12823-VC), Gata6 (Thermo Fisher Scientific, VB1-15776-VC), Zscan4d (Thermo Fisher Scientific, VB4-3138327-VC), Zscan4f (Thermo Fisher Scientific, VB4-3139209-VC).

### 10× Genomics

ES knockdown cells were collected greater than 5×10^6 by FACS. FACS-enriched and total single cell were loaded for droplet-based scRNA-seq according to the manufacturer’s protocol for the Chromium Single Cell 5′ gene expression v.2 (10x Genomics) to obtain 8,000–10,000 cells per reaction. Library preparation was carried out according to the manufacturer’s protocol. 2 ibraries were sequenced across both lanes of an Illumina NovaSeq 6000 S2 flow cell with 150 bp paired-end reads.

### Co-immunoprecipitation (Co-IP) assays

Co-immunoprecipitation (Co-IP) assays were performed on whole cell lysate of WT and KDL1. Cell pellets were resuspended in Cell lysis buffer (Beyotime, P0013) containing protease inhibitors (Beyotime, P1045) incubated for 20 min on ice, and centrifuged 12000g 10min at 4℃, then transfer cell suspensions to new tube. Supernatants containing target protein were quantified using a BCA protein quantification kit (Yeasen, 20201ES76), and 200ug protein extracts were immunoprecipitated using the primary antibodies NANOG (Abcam, ab203919), rotating overnight at 4℃. The following day, immune complexes were bound to 20ul ProteinA/G Dynabeads (Thermo Fisher Scientific), washed 5 times in cell lysis buffer at 4℃ , boiled in 5×Loading Buffer (Epizyme, LT102S) and used for western blotting analysis.

### Western blotting

These samples were prepared from CoIP to perform western blotting. Separation gels using PAGE Gel Fast Preparation kit (Epizyme, PG112) and then transferred to PVDF membranes. Blocking was performed for 45 min in 5% milk/PBS-T buffer followed by incubation overnight shake softly with primary antibodies (EZH2, CST; NANOG) at 4℃. The following day, membranes were incubated with the appropriate anti-mouse/rabbit secondary antibodies conjugated to HRP (Jackson) for 1h, and proteins were detected by ECL or ECL Plus reagent and autoradiography. To assess the protein expression levels of TET1, cells exhibiting different pluripotency were harvested and subjected to protein extraction using RIPA Buffer (P0013B; Beyotime). Subsequently, western blotting (WB) was conducted using the TET1 primary antibodies (GTX124207, GeneTex).

### MARGI experiment

The MARGI experiment in E14TG2a ESCs was performed as previously described (Sridhar et al. 2017). Briefly, the 4-5 million cells were crosslinked by 1% formaldehyde (Thermo, #28906). Chromatin was fragment using HaeIII (NEB, #R0108M) and subsequently stabilized on streptavidin T1 beads (Thermo, #65601) after cell lysis and biotinylation of proteins (Thermo, #21334). Then, RNA and DNA ends were prepared for ligation. First, a non-templated dA tail was added using Exo(-) Klenow (NEB, #M0212S) to 3’-P presenting in the RNA that was dephosphorylated using T4 PNK (NEB, #M0201S). Linker and RNA ligation was performed by ligating pre-adenylated linker to the 3’-OH of RNA using RNA ligase2, truncated KQ (NEB, #M0373L). Reaction was carried out at 22℃ for 6 hours followed by 16℃ overnight. Second, DNA ends were phosphorylated using T4 PNK (NEB, #M0201S) and T4 DNA ligase (NEB, #M0202M) were used for proximity ligation by 16℃ overnight. After reverse crosslinking and DNA/RNA extraction, the RNA was reverse transcribed using Supercript IV (Invitrogen, #18090050) and purified the single strand DNA without biotin using Silane beads (Invitrogen, #37002D). Then, purifying DNA was circularized and digested by restriction enzyme BamHI (NEB, #R3136S). Finally, the right and left flanking sequences of the linker were split as primers for amplification and sequencing.

### Detection for distal transposon RNA-chromatin interactions from MARGI data

1. Alignment of DNA and RNA reads: The two ends of RNA-DNA paired reads were separately processed. Firstly, reads processing, including removing adaptor, filtering low-quality and duplication reads, was performed according to previously published pipeline e (Sridhar et al. 2017). Then, we ran STAR (Dobin et al. 2013) using the following options: --runThread N – genomeDir—alignEndsType Extend5pOfRead1 --seedSearchStartLmax 25 --outFilterScoreMin 21--outFilterScoreMinOverLread 0.13 --outFilterMatchNmin 21 -- outFilterMatchNminOverLread 0.13 --alignIntronMax 1 and BWA (Li and Durbin 2009) MEM using the following options: -k 15 –T 15 for DNA and RNA ends, respectively.
2. The RNA ends of RNA-DNA paired reads were mapped to transposon element regions that were annotated in UCSC RepeatMasker.
3. Mappable DNA reads with multiple hits were removed.
4. We filtered the mappable DNA reads that were annotated in heterochromatin regions via H3K9me3 and HP1a ChIP-seq data (GSE57092, GSE39579, ENCODE). Exceptionally the LINE-1 DNAs in heterochromatin regions co-bound by LINE-1 RNA-KAP1 were retained.
5. The definition of the term “distal” followed the rule that the chromosomal coordinates between the mappable RNA end and DNA end of RNA-DNA paired reads should be more than 5kb apart.
6. The DNA loci targeted by transposon RNAs, spanning ∼200bp region, are defined by the clusters of distal RNA-DNA paired reads.

### RNA-DNA hybrid prediction between the sequence contexts of a transposon RNA-chromatin interaction pair

10,000 DNA sequences outside the transposon RNA targeting regions were randomly sampled for 100 times to compose the pseudo transposon RNA-DNA pairs as control. And 20,000 confident distal interaction pairs between non-transposon RNAs and chromatins were selected to serve as another control group. The repeat information was annotated by UCSC RepeatMasker. The perfect-matched sequence pairs form the positives. Predictions of triple helix and DNA/RNA hybrid duplex structure were performed using Triplexator and RNAhybrid (Kruger and Rehmsmeier 2006; Buske et al. 2012) on the above datasets. A 50bp sliding window along transposon RNA was utilized for RNAhybrid prediction. The minimal free energy defines the binding strength of RNA-DNA hybrid duplex.

### Peak calling from R-loop and histone modification/protein ChIP-seq data

We re-analyzed R-loop data were downloaded from NCBI (GSE145964, GSE70189). The raw data processing was the same as pxMARGI data and uniquely aligned to hg19 or mm10 genome by STAR (Dobin et al. 2013) with default parameters. MACS2 (Zhang et al. 2008) with the --broad parameter was used for peak calling. H3K9me3 stable peaks in H9 (ENCODE) and H2A119KUb1 peaks in mESC (GSE145964) were used, directly. H3K27me3 (ENCODE, Epigenome), H3K9me3 (ENCODE, GSE57092) and HP1a (GSE57092, GSE39579) ChIP-seq data of the above mentioned were downloaded. For raw FASTQ data, we used Bowtie2 (Langmead and Salzberg 2012) or BWA (Li and Durbin 2009) with the default parameters for alignment of ChIP-seq data. MACS2 (Zhang et al. 2008) was used for peak calling from BAM or BED files.

### RNA-seq data processing

Public available FASTQ files from H9, HEK293 and E14 RNA-seq experiments from the Gene Expression Omnibus (GSE103715, GSE43572, GSE54106) were removed PCR duplicated reads and ran BWA (Li and Durbin 2009)MEM using the same options as the RNA ends of read pairs from MARGI experiments to reduce technical bias. For Kap1-KO RNA-seq data (GSE99215) in H9, reads processing was performed using the same options and mapped to mm10 or hg19 genome using STAR (Dobin et al. 2013) with the following parameters: -- outFilterMultimapNmax 100 --outFilterMismatchNoverReadLmax 0.04 --alignIntronMax 1000000 --alignMatesGapMax 1000000, that enabled to output multiple alignments for a read. Normalized reads count per gene in Kap1-KO experiment were calculated based on the total mappable reads of Kap1-KO and WT data. For PRC-KO RNA-seq data (GSE66814, GSE132753), the results of differential expression analysis were used, directly.

### Genomic annotations on aligned reads or called peaks

Genomic information, including promoter (+5/-1kb from TSS), CDS, UTR, intron and intergenic regions, and gene symbols, was extracted from Ensembl (Homo sapiens.GRCh37.75, Mus musculus.GRCm38.79). The aligned reads and called peaks from high-throughput sequencing data were annotated with the genomic information. The biomaRt package (Durinck et al. 2009) was used for the gene symbol conversion between human and mouse species.

### The definition of co-localization between two binding loci

The co-localization of binding sites of protein/histone modifications or R-loop regions was determined via BEDTools (Quinlan and Hall 2010) if these regions located within 500bp region of transposon RNA binding loci. The binding sites of transcription factors downloaded from GTRD (Gene Transcription Regulation Database) (Yevshin et al. 2017), that is the most complete collection of uniformly processed ChIP-seq data on identification of transcription factor binding sites for human and mouse.

### Gene Ontology and Pathway Enrichment Analysis

Gene Ontology enrichment analysis was conducted by David.

### Quantification and statistical analysis

All statistical analyses were performed in R/Bioconductor. All statistics were * p < 0.05, ** p < 0.01, *** p < 0.001, **** p < 0.0001. They were calculated by Wilcox test unless specially noted otherwise.

### Spent blastocyst medium collection

A total of 10 μl blastocyst medium (BM) from each embryo was loaded into RNase-DNase-free PCR tubes, when the embryos reached a fully expanded blastocyst stage, generally between day 5 or day 6 of embryos cultured, and immediately stored at –80◦C until further experiment.

### Library preparation and sequencing

The RNA-seq library of BM using a previously described method SMART-seq2 (Picelli et al. 2014) and modified some procedures to prepare. Briefly, 10 μl BM was transferred into a PCR tube, and add 1 μl protein K was incubated at 50◦C for 1.5h, 75℃ for 30min. Ten cycles were applied for the connection of Illumina Adapters and Indexes finally. Purification of final RNA-seq library was processed using VAHTS@ DNA Clean Beads (Vazyme Biotech Co., Ltd). The cDNA library concentration was measured using a Qubit® 2.0 fluorometer. The library was sequenced by Illumina HiSeq 2500 (Novogene, Tianjin) sequencer using a 2 × 150 bp paired-end pattern (PE150).

## Acknowledgments

We thank professor Yungui Yang for constructive criticism and fruitful discussion. We thank Dr. Minglei Shi, Dr. Yang Chen and Professor Michael Zhang for kind help on experiment protocol.

This work was supported by grants from the National Key Research and Development Program of China (2021YFF1200904 and 2024YFF0729204 to J.C.), the Beijing Natural Science Foundation (Z210011 to J.C.), and the National Natural Science Foundation of China (32070795 and 32471512 to J.C.).

## Author Contributions

J.C. and C.Z. envisioned the project, funding acquisition. L.R., W.X., M.L. and F.S. implemented the experiment. Y.T. and J.H. performed the data analysis. J.C., W.X., L.R and J.R. wrote the paper. C.Z. provided assistance in writing and analysis.

## Reference

Bell JC, Jukam D, Teran NA, Risca VI, Smith OK, Johnson WL, Skotheim JM, Greenleaf WJ, Straight AF. 2018. Chromatin-associated RNA sequencing (ChAR-seq) maps genome-wide RNA-to-DNA contacts. Elife 7.

Beltran M, Yates CM, Skalska L, Dawson M, Reis FP, Viiri K, Fisher CL, Sibley CR, Foster BM, Bartke T et al. 2016. The interaction of PRC2 with RNA or chromatin is mutually antagonistic. Genome Res 26: 896–907.

Bonetti A, Agostini F, Suzuki AM, Hashimoto K, Pascarella G, Gimenez J, Roos L, Nash AJ, Ghilotti M, Cameron CJF et al. 2020. RADICL-seq identifies general and cell type-specific principles of genome-wide RNA-chromatin interactions. Nat Commun 11: 1018.

Bourque G, Leong B, Vega VB, Chen X, Lee YL, Srinivasan KG, Chew JL, Ruan Y, Wei CL, Ng HH et al. 2008. Evolution of the mammalian transcription factor binding repertoire via transposable elements. Genome Res 18: 1752–1762.

Bulut-Karslioglu A, De La Rosa-Velazquez IA, Ramirez F, Barenboim M, Onishi-Seebacher M, Arand J, Galan C, Winter GE, Engist B, Gerle B et al. 2014. Suv39h-dependent H3K9me3 marks intact retrotransposons and silences LINE elements in mouse embryonic stem cells. Mol Cell 55: 277–290.

Castro-Diaz N, Ecco G, Coluccio A, Kapopoulou A, Yazdanpanah B, Friedli M, Duc J, Jang SM, Turelli P, Trono D. 2014. Evolutionally dynamic L1 regulation in embryonic stem cells. Genes Dev 28: 1397–1409.

Chen C, Liu W, Guo J, Liu Y, Liu X, Liu J, Dou X, Le R, Huang Y, Li C et al. 2021. Nuclear m(6)A reader YTHDC1 regulates the scaffold function of LINE1 RNA in mouse ESCs and early embryos. Protein Cell 12: 455–474.

Cifuentes-Rojas C, Hernandez AJ, Sarma K, Lee JT. 2014. Regulatory interactions between RNA and polycomb repressive complex 2. Mol Cell 55: 171–185.

De Iaco A, Planet E, Coluccio A, Verp S, Duc J, Trono D. 2017. DUX-family transcription factors regulate zygotic genome activation in placental mammals. Nat Genet 49: 941–945.

Deniz O, Frost JM, Branco MR. 2019. Regulation of transposable elements by DNA modifications. Nat Rev Genet 20: 417–431.

Di Croce L, Helin K. 2013. Transcriptional regulation by Polycomb group proteins. Nat Struct Mol Biol 20: 1147–1155.

Ding N, You A, Zhao S, Yang H, Lai C, Ye F. 2023. EZH2 inhibitor Tazemetostat synergizes with JQ-1 in esophageal cancer by inhibiting c-Myc signaling pathway. Med Oncol 40: 281.

Fadloun A, Le Gras S, Jost B, Ziegler-Birling C, Takahashi H, Gorab E, Carninci P, Torres-Padilla ME. 2013. Chromatin signatures and retrotransposon profiling in mouse embryos reveal regulation of LINE-1 by RNA. Nat Struct Mol Biol 20: 332–338.

Fu X, Djekidel MN, Zhang Y. 2020. A transcriptional roadmap for 2C-like-to-pluripotent state transition. Sci Adv 6: eaay5181.

Giordano J, Ge Y, Gelfand Y, Abrusan G, Benson G, Warburton PE. 2007. Evolutionary history of mammalian transposons determined by genome-wide defragmentation. PLoS Comput Biol 3: e137.

Hao Y, Wang D, Wu S, Li X, Shao C, Zhang P, Chen JY, Lim DH, Fu XD, Chen R et al. 2020. Active retrotransposons help maintain pericentromeric heterochromatin required for faithful cell division. Genome Res 30: 1570–1582.

Imbeault M, Helleboid PY, Trono D. 2017. KRAB zinc-finger proteins contribute to the evolution of gene regulatory networks. Nature 543: 550–554.

Jachowicz JW, Bing X, Pontabry J, Boskovic A, Rando OJ, Torres-Padilla ME. 2017. LINE-1 activation after fertilization regulates global chromatin accessibility in the early mouse embryo. Nat Genet 49: 1502–1510.

Johnson R, Guigo R. 2014. The RIDL hypothesis: transposable elements as functional domains of long noncoding RNAs. RNA 20: 959–976.

Karaky M, Fedetz M, Potenciano V, Andres-Leon E, Codina AE, Barrionuevo C, Alcina A, Matesanz F. 2018. SP140 regulates the expression of immune-related genes associated with multiple sclerosis and other autoimmune diseases by NF-kappaB inhibition. Hum Mol Genet 27: 4012–4023.

Kashef J, Franz CM. 2015. Quantitative methods for analyzing cell-cell adhesion in development. Dev Biol 401: 165–174.

Kruger J, Rehmsmeier M. 2006. RNAhybrid: microRNA target prediction easy, fast and flexible. Nucleic Acids Res 34: W451–454.

Kunarso G, Chia NY, Jeyakani J, Hwang C, Lu X, Chan YS, Ng HH, Bourque G. 2010. Transposable elements have rewired the core regulatory network of human embryonic stem cells. Nat Genet 42: 631–634.

Lander ES, Linton LM, Birren B, Nusbaum C, Zody MC, Baldwin J, Devon K, Dewar K, Doyle M, FitzHugh W et al. 2001. Initial sequencing and analysis of the human genome. Nature 409: 860–921.

Li H, Durbin R. 2009. Fast and accurate short read alignment with Burrows-Wheeler transform. Bioinformatics 25: 1754–1760.

Li X, Bie L, Wang Y, Hong Y, Zhou Z, Fan Y, Yan X, Tao Y, Huang C, Zhang Y et al. 2024. LINE-1 transcription activates long-range gene expression. Nat Genet 56: 1494–1502.

Li X, Fu XD. 2019. Chromatin-associated RNAs as facilitators of functional genomic interactions. Nat Rev Genet 20: 503–519.

Li X, Zhou B, Chen L, Gou LT, Li H, Fu XD. 2017. GRID-seq reveals the global RNA-chromatin interactome. Nat Biotechnol 35: 940–950.

Lin YC, Boone M, Meuris L, Lemmens I, Van Roy N, Soete A, Reumers J, Moisse M, Plaisance S, Drmanac R et al. 2014. Genome dynamics of the human embryonic kidney 293 lineage in response to cell biology manipulations. Nat Commun 5: 4767.

Liu J, Gao M, He J, Wu K, Lin S, Jin L, Chen Y, Liu H, Shi J, Wang X et al. 2021. The RNA m(6)A reader YTHDC1 silences retrotransposons and guards ES cell identity. Nature 591: 322–326.

Low Y, Tan DEK, Hu Z, Tan SYX, Tee WW. 2021. Transposable Element Dynamics and Regulation during Zygotic Genome Activation in Mammalian Embryos and Embryonic Stem Cell Model Systems. Stem Cells Int 2021: 1624669.

Lu JY, Chang L, Li T, Wang T, Yin Y, Zhan G, Han X, Zhang K, Tao Y, Percharde M et al. 2021. Homotypic clustering of L1 and B1/Alu repeats compartmentalizes the 3D genome. Cell Res 31: 613–630.

Mangoni D, Simi A, Lau P, Armaos A, Ansaloni F, Codino A, Damiani D, Floreani L, Di Carlo V, Vozzi D et al. 2023. LINE-1 regulates cortical development by acting as long non-coding RNAs. Nat Commun 14: 4974.

McCallie BR, Parks JC, Trahan GD, Jones KL, Coate BD, Griffin DK, Schoolcraft WB, Katz-Jaffe MG. 2019. Compromised global embryonic transcriptome associated with advanced maternal age. J Assist Reprod Genet 36: 915–924.

Meng S, Liu X, Zhu S, Xie P, Fang H, Pan Q, Fang K, Li F, Zhang J, Che Z et al. 2023. Young LINE-1 transposon 5’ UTRs marked by elongation factor ELL3 function as enhancers to regulate naive pluripotency in embryonic stem cells. Nat Cell Biol 25: 1319–1331.

Mita P, Boeke JD. 2016. How retrotransposons shape genome regulation. Curr Opin Genet Dev 37: 90–100.

Mouse Genome Sequencing C, Waterston RH, Lindblad-Toh K, Birney E, Rogers J, Abril JF, Agarwal P, Agarwala R, Ainscough R, Alexandersson M et al. 2002. Initial sequencing and comparative analysis of the mouse genome. Nature 420: 520–562.

Percharde M, Lin CJ, Ramalho-Santos M. 2021. Depletion of nuclear LINE1 RNA in mouse ESCs and embryos. STAR Protoc 2: 100726.

Percharde M, Lin CJ, Yin Y, Guan J, Peixoto GA, Bulut-Karslioglu A, Biechele S, Huang B, Shen X, Ramalho-Santos M. 2018. A LINE1-Nucleolin Partnership Regulates Early Development and ESC Identity. Cell 174: 391–405 e319.

Pezic D, Manakov SA, Sachidanandam R, Aravin AA. 2014. piRNA pathway targets active LINE1 elements to establish the repressive H3K9me3 mark in germ cells. Genes Dev 28: 1410–1428.

Picelli S, Faridani OR, Björklund ÅK, Winberg G, Sagasser S, Sandberg R. 2014. Full-length RNA-seq from single cells using Smart-seq2. Nat Protoc 9: 171–181.

Pijuan-Sala B, Griffiths JA, Guibentif C, Hiscock TW, Jawaid W, Calero-Nieto FJ, Mulas C, Ibarra-Soria X, Tyser RCV, Ho DLL et al. 2019. A single-cell molecular map of mouse gastrulation and early organogenesis. Nature 566: 490–495.

Pizarro JG, Cristofari G. 2016. Post-Transcriptional Control of LINE-1 Retrotransposition by Cellular Host Factors in Somatic Cells. Front Cell Dev Biol 4: 14.

Sanz LA, Hartono SR, Lim YW, Steyaert S, Rajpurkar A, Ginno PA, Xu X, Chedin F. 2016. Prevalent, Dynamic, and Conserved R-Loop Structures Associate with Specific Epigenomic Signatures in Mammals. Mol Cell 63: 167–178.

Scialdone A, Tanaka Y, Jawaid W, Moignard V, Wilson NK, Macaulay IC, Marioni JC, Gottgens B. 2016. Resolving early mesoderm diversification through single-cell expression profiling. Nature 535: 289–293.

Sridhar B, Rivas-Astroza M, Nguyen TC, Chen W, Yan Z, Cao X, Hebert L, Zhong S. 2017. Systematic Mapping of RNA-Chromatin Interactions In Vivo. Curr Biol 27: 610–612.

Sundaram V, Cheng Y, Ma Z, Li D, Xing X, Edge P, Snyder MP, Wang T. 2014. Widespread contribution of transposable elements to the innovation of gene regulatory networks. Genome Res 24: 1963–1976.

Testori A, Caizzi L, Cutrupi S, Friard O, De Bortoli M, Cora D, Caselle M. 2012. The role of Transposable Elements in shaping the combinatorial interaction of Transcription Factors. BMC Genomics 13: 400.

Wang R, Zhang P, Wang J, Ma L, E W, Suo S, Jiang M, Li J, Chen H, Sun H et al. 2023. Construction of a cross-species cell landscape at single-cell level. Nucleic Acids Res 51: 501–516.

Wang X, Goodrich KJ, Gooding AR, Naeem H, Archer S, Paucek RD, Youmans DT, Cech TR, Davidovich C. 2017. Targeting of Polycomb Repressive Complex 2 to RNA by Short Repeats of Consecutive Guanines. Mol Cell 65: 1056–1067 e1055.

Warkocki Z, Krawczyk PS, Adamska D, Bijata K, Garcia-Perez JL, Dziembowski A. 2018. Uridylation by TUT4/7 Restricts Retrotransposition of Human LINE-1s. Cell 174: 1537–1548 e1529.

